# Experimental comparison of mowing and mulching on snail communities in wet meadows

**DOI:** 10.1101/2024.11.16.623914

**Authors:** Roland Farkas, Miklós Bán, Gergő Oláh, György Dudás, Zoltán Barta

## Abstract

The condition of wet meadows nowadays depends mainly on human activities; the biodiversity and productivity of these habitats can only be maintained by appropriate management methods. Mowing and grazing are well-known traditional methods, but a new method, regular mulching – the fragmentation of plant material and its deposition in the area - is also increasingly used. The ecological impact of traditional management is still not fully known, and we know almost nothing about the effects of mulching. To better understand the impact of these methods we directly compared their effects on snails, an invertebrate taxon common in these habitats and known as good bioindicators. We experimentally manipulated management methods on two wet meadows in Northern Hungary, Europe, and surveyed their snail communities immediately before and one year after treatments. Our results showed that mowing had a detectable negative effect on the snail communities. On the other hand, mulching did not alter their characteristics. Therefore, mulching may be a more desirable method for short-term interventions on wet meadows, whilst at the same time also preserving their biodiversity.

## Introduction

Grasslands are an essential habitat for many animals. They make up 40.5 percent of land habitats worldwide (Petermann and Buzhdygan, 2021). Many of the species that live here are specialist or endemic. In addition to their high ecological value, they are also essential economically. Indeed, human culture developed on or near grasslands, which were necessary for, first the nomadic lifestyle of livestock grazing, and then the static, large-scale rearing of livestock and growing crops. (Squires et al 2018). The extent and condition of grasslands have changed over time; since the agricultural revolution, this has been largely in response to human needs and activities (Poschlod and WallisDeVries 2002). While a few activities, such as deforestation, even create new grasslands, in general their extent has decreased at an ever-faster rate with the spread of intensive agriculture and urbanisation. The constant need for increasing efficiency, and hence the emergence of new agricultural technologies, are strongly shaping these habitats (Fischer et al 2008). Today, the vast majority of historical grasslands have been transformed and cultivated by humans, for rearing and grazing farm animals (Hejcman et al 2013) and only small areas of pristine grassland remain. Many species of historical grasslands were able to settle and survive on cultivated grasslands before the original habitats had disappeared. The maintenance of today’s secondary grasslands is therefore a priority task. Thus the fate of grasslands is closely tied to humanity and their survival strongly depends on their proper management by humans.

Management practices are shaped by social changes, political and legal regulations, directives and agri-environmental programmes that transcend national borders, and thus are dictated by many stakeholders at various levels (Stoate et al. 2009). In recent times, fewer people are benefiting from grasslands. The ownership of grasslands has become more centralised, with individual owners farming ever-larger areas and their agriculture is largely characterised by more intensive farming on these larger plots. (Marini et al. 2011). The adverse impact of these large-scale changes on biodiversity is shown by the negative correlation found between the species richness of a grassland and the amount of money spent on farming it (Zechmeister et al. 2003).

A specific but widespread form of grasslands are the wet meadow, defined as areas where the soil is saturated with water in the whole year or at least a part of the growing season. Because they occur only under specific hydrological conditions, wet meadows are highly sensitive to environmental changes. However, these hydrological conditions are negatively affected by both current climatic changes and farming (Joyce et al. 2016). As a result, the extent of wet meadows is decreasing in all EU countries and they are classified as threatened by the European Red List of Habitats (Jansen et al. 2016).

The current thinking is that in general, it is desirable to maintain grasslands, rather than allowing them to return to the ancestral forested state, both because they are important for farming, but also from a conservation perspective because of their high biodiversity (Petermann and Buzhdygan, 2021). Grazing and mowing are traditional agricultural activities that are able to prevent spontaneous reforestation. Grazing is the process whereby the animals eat the upper part of the vegetation, thus removing it from the area. In mowing, the vegetation is cut off near the surface and, after appropriate treatment, removed from the area. In the absence of these activities, natural succession processes usually take place. The emergence of shrub and woody vegetation hinders farming (Moog et al. 2002) and threatens the continued existence of grasslands. A striking example of these processes is Central and Eastern Europe, where the political changes of the late 1980’s and early 1990’s drastically altered the agricultural landscape. Large-scale changes in ownership and economic restructuring led to the regional abandonment of large areas, many of which are still in this state today (Kuemmerle et al. 2008, Halada et al. 2017).

The process of mechanised mulching has been developed to manage (Mašková et al 2009) and recultivate abandoned grassland habitats (Liira et al 2009). Despite the fact that mulching has several different meanings in English, the specific mulching method is often not clearly defined in scientific articles (e.g. Pižl and Starý 2001, Babálová and Štrbová 2014). This methodological ambiguity makes it difficult to understand the effects of mulching on biodiversity. One common meaning of mulching is the process of spreading plant material that has been harvested from another location, for example to protect economic crops (e.g. Bautista 2018). In other cases, it is used to mean when plant material cut in an area by mowing is not collected but left to decompose on the spot (e.g. Liira et al 2009, Klimeš et al. 2013). In this article, we use a third common definition of mulching, whereby the standing vegetation on the lawn is mowed then shredded into small pieces, with the resulting material spread evenly over the surface of the meadow (and not removed). This definition is also used by Moog et al (2002), Schmitt (2003), Doležal et al. (2011), Pavlů et al. (2016), Gaisler et al (2019), and Georgi et al (2022, 2023). Regular mulching can be used to remove shrub and smaller trees from abandoned grasslands and restore an area in a matter of a few years. It is commonly used for habitat restoration in nature conservation areas. One of the reasons for the spread of mulching is that it is more economical than traditional methods (Liira et al. 2009). Mowing and mulching can have various effects on the environment. Direct effects from the moving parts of the machines cause damage to the flora and fauna (Humbert et al. 2010, Steidle et al. 2022). Indirect effects are when the physical and chemical properties of the habitat are changed (Humbert et al. 2009). In this case, it is important whether the plant material is removed from the area or left to decompose in place. As vegetation is removed, direct irradiance penetrates deeper, even to the surface of the soil. This can result in less favourable temperatures compared to similar habitats with closed vegetation (Gardiner and Hassal 2009, Hesslerová et al. 2013). At the same time, covering the surface with cut plant material results in lower temperatures (Dudas et al. 2016) and higher soil humidity (Shirish et al. 2013). Mowing and mulching also changes the soil microbiome, influencing decomposition processes and the amount of available nutrients (Kaštovská et al. 2005, Pavlů et al. 2016, Horan et al. 2018). Organic matter accumulated on the soil surface significantly increases the activity of decomposing organisms and enzymes in the soil (Jabran 2022). It is important to understand the overall effect of these many factors in order to improve management practices.

The impact of management on grasslands depends greatly on the method used and the characteristics of the habitat concerned, and is an important research topic. Most work so far has looked at how management practices change plant communities, in particular their species composition, biomass or priority species groups. Plant communities in mowed areas have generally higher diversity than those in unmanaged, abandoned areas, except when the area is intensively farmed. This has especially been demonstrated in dry, semi-arid, nutrient-poor meadows and mountain pastures (Ryser et al. 1995). In these areas, both mowing and mulching have generally been found to have similar positive results (Zelený et al. 2001, Moog et al. 2002, Gaisler et al. 2004, Klimeš et al. 2013, Doležal et al. 2011). However, wet meadows, a more nutrient-rich habitat, are significantly less researched. What little evidence we have indicates, that mulching in wet, alluvial habitats may aid habitat restoration (Liira et al. 2009).

In contrast to our more detailed knowledge of the impact of mowing and mulching on flora, their effects on fauna, especially on invertebrates, have received much less focus. According to the current literature, mowing mostly has a negative impact on some invertebrate groups (snails: Pech et al. 2015, Farkas et al. 2024, spiders: Cattin et al. 2003, dung beetles: Frank et al. 2017, orthopterans: Chisté et al. 2016). Mulching has also been shown to have a similar negative effect (butterflies: Babálová and Štrbová 2014, bees and wasps: Georgi et al. 2022, insect larvae, flower visitors: Georgi et al. 2023). However, the impact of these management methods can be influenced by a number of factors, such as the time of year and the frequency of treatments (Humbert et al. 2009, Georgi et al. 2023). The literature suggests that it is difficult to draw a general conclusion about the effect of grassland management on animals because different taxa living in the same habitat have been observed to respond differently to the same treatment (Pižl and Starý 2001, Cizek et al. 2012).

Snails (Gastropoda) are common and widespread inhabitants of wet meadows (Welter-Schultes 2012). They are significant actors in the decomposition process (Newell, 1967). They stimulate the activity of microbial detritivores by fragmenting organic materials, and their excretion of feces and mucus production provides favorable conditions for microbial life (Theenhaus and Scheu, 1996). Furthermore, the high water content of their bodies, their slow movement and limited dispersal ability make snail populations especially sensitive to environmental change. This makes them excellent subjects for studying the effects of environmental factors in general (Pech et al. 2015, Wehner et al. 2019). Nevertheless, the impact of mulching on snail communities is largely unknown. At most, conclusions can be drawn from experience with one kind of mulching when crop fields are covered with cut plant matter (Bautista et al. 2008) - but crop fields and grasslands are fairly different habitats.

The aim of our study was to identify sustainable management methods for wet meadows by using snail populations as indicator species. We also wanted to know whether it is possible to say after the first year that a treatment is harmful to the snail community. We selected undisturbed wet meadows that had remained untouched for decades. We then characterized snail communities within these habitats. Using commonly utilized agricultural machinery, we mowed and mulched portions of the areas according to a stratified experimental design. After one year, we returned to collect data on the current state of the snail communities and investigated the effects of each treatment on them.

## Methods

### Study area

The experiment was carried out at two sites in Northern Hungary, Europe (**Fig 1**). Both of them are located in stream valleys of a hilly area, west of the Bükk Mountains. This area is characterised by high forest cover, with wet meadows occurring almost exclusively along the valley bottom watercourses. Annual rainfall in this area ranges between 600-700 mm and the average annual temperature is around 9℃ (Dövényi 2010). In the valley bottoms meadow moulding soils have developed on clayey, loamy, sandstone bedrock (Dövényi 2010). In both sites, the habitats studied were non-tussock tall-sedge beds, with a dominance of sedge (*Carex acutiformis*). The characteristic associated species observed were *Lysimachia vulgaris*, *Lythrum salicaria*, *Cirsium oleraceum* and *Urtica dioica*. Both sites were flat, horizontal areas without any obvious microhabitat structure. No bushes or trees were present in any of the sites. An important criterion for the selection of the sites was that they represent wet meadows that have not been farmed for the last 20-30 years. To determine this, a series of archive aerial photographs (www.fentrol.hu) were used.

**Figure 1.**
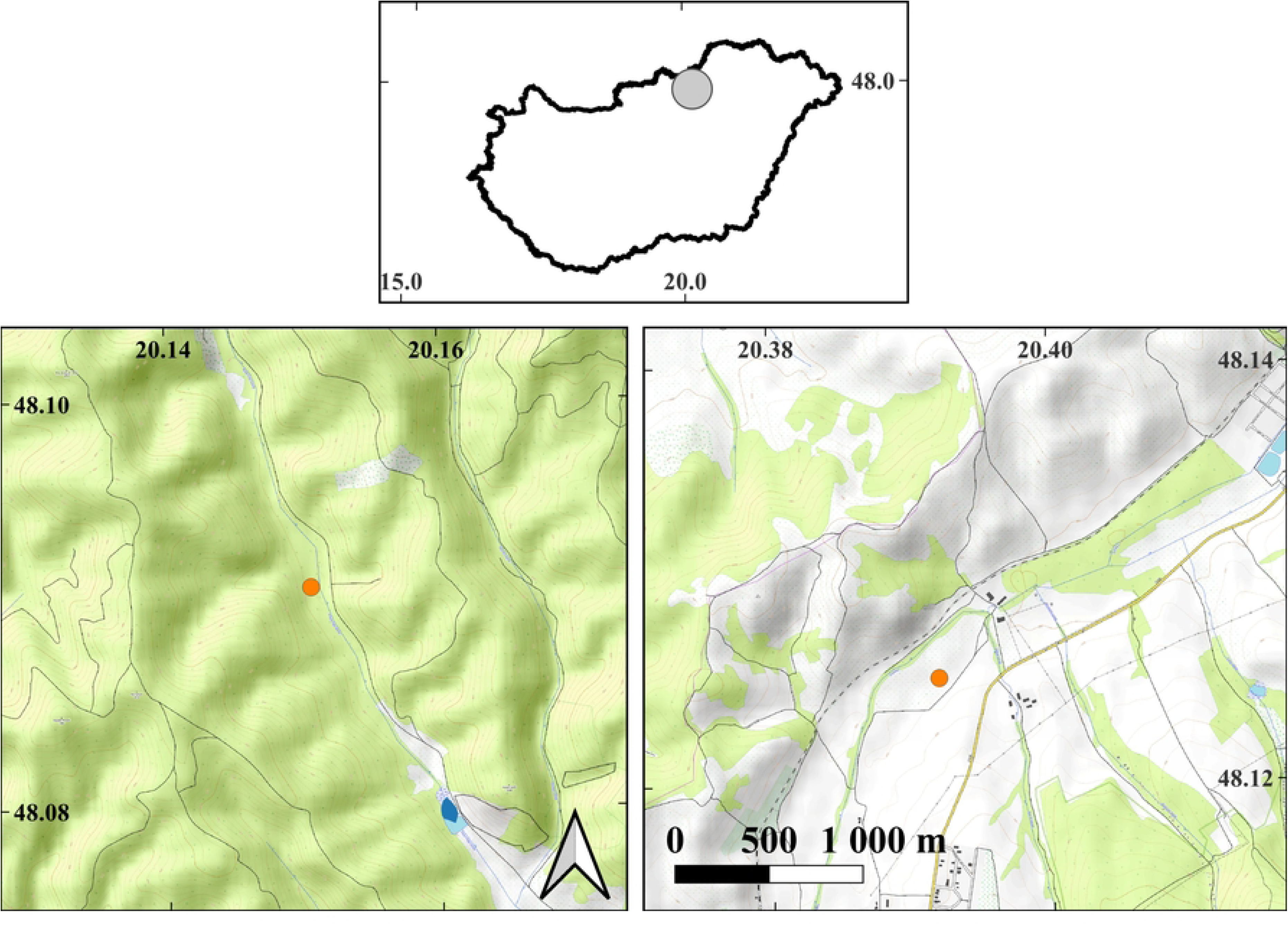
The map of the study sites. Top center: the borders of Hungary are visible, and a gray circle indicates the location of the study areas. In the bottom row, detailed maps depicting the locations of sample sites (orange dots) are presented. Left side: Tarnalelesz sample site, right side: Szilvásvárad sample site. Both detailed maps share the same scale and orientation. WGS 84 geographic coordinates are marked on the axes. Map data: © OpenStreetMap contributors, SRTM | Map display: © OpenTopoMap (CC-BY-SA).

### Experimental design

A sample area of 45×15 metres was marked out in each site. These were divided into 9 parallel parcels of 5 x 15 metres. Both the edges of the sample area and the boundaries of the parcels were marked with permanent stakes. Out of the 9 parcels, 3 were mowed, 3 were mulched, and 3 remained untouched as control plots. Treatments were assigned to the parcels in a stratified random way: each block of three consecutive parcels received all three treatments in a random order. In every parcel, one sampling point was assigned to each 5×3 meter plot, resulting in 45 sampling points at each site (**Fig 2**)

**Figure 2.**
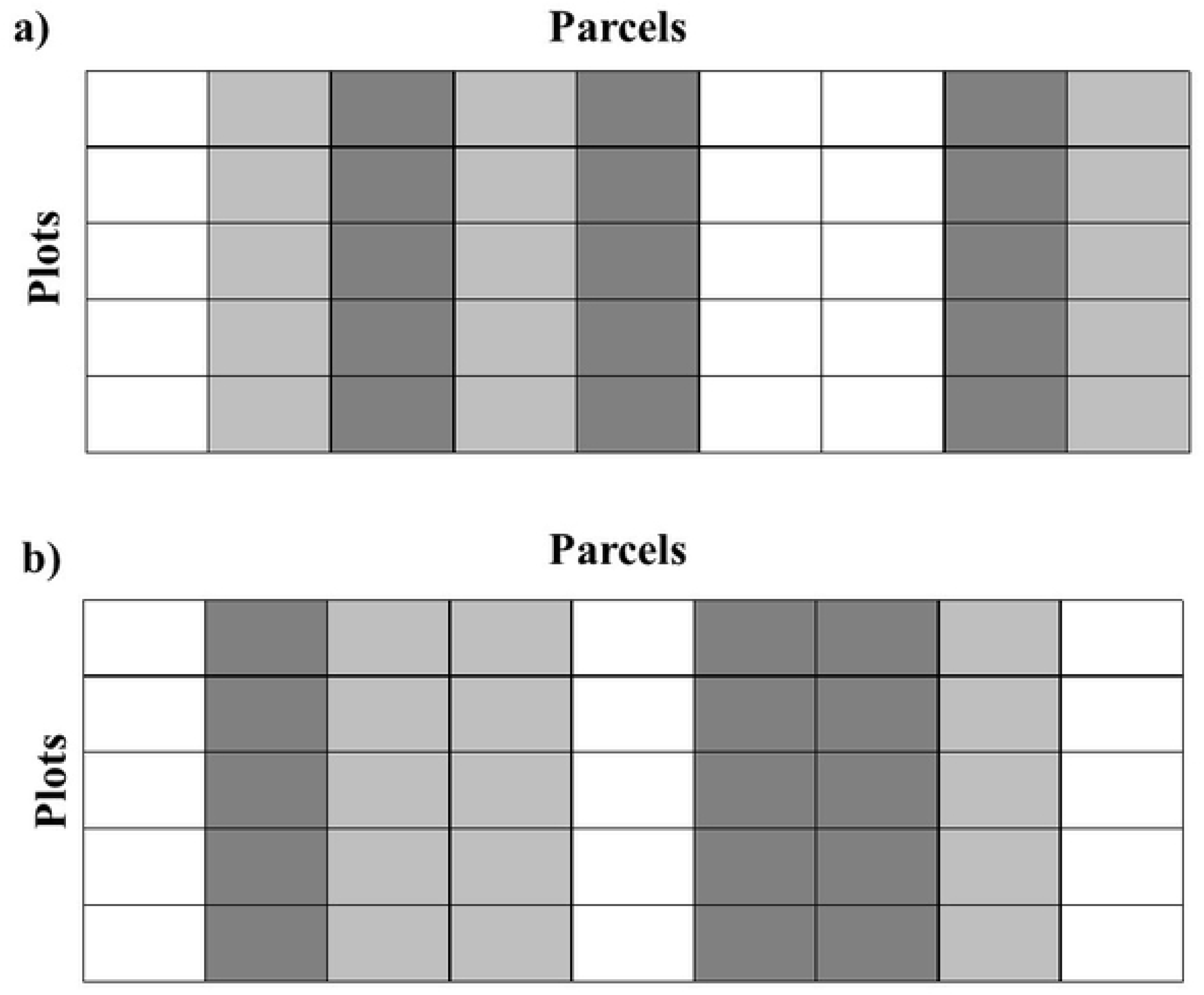
Detailed sampling design. a) Szilvásvárad site, b) Tarnalelesz site. The columns represent treatment parcels, and the rows indicate sampling plots. Different colors indicate the type of treatment: white – control, light gray – mowed, dark gray – mulched.

On June 28, 2021. (Szilvásvárad) and June 29, 2021. (Tarnalelesz), a malacological sampling was taken in each plot before the start of management. Treatments were carried out once a few days after this sampling, on July 1, 2021 at both sites. This is a typical time for farmers in the region to carry out farming activities on meadows. In the mowed parcels, we cut the vegetation with a mower machine pulled by a tractor. The cut material was immediately removed from the surface using hand rakes. Mulching treatment was done by a mulching machine pulled by a tractor to shred the vegetation into pieces smaller than 10 cm. The resulting material was not removed and evenly covered the surface. Both sites were then left undisturbed for approximately one year. Subsequently, on August 15, 2022 (Szilvásvárad) and August 14, 2022 (Tarnalelesz), we went back to the sites and again collected a malacological sample from each plot. Malacological sampling was carried out in the morning hours each time. Sampling dates were selected to ensure that the weather conditions were similar and rain-free during the two-day periods.

One malacological sample was taken per plot per sampling occasion. Sampling was performed by collecting the leaf litter and the top 1 cm thick layer of the soil from a 25X25 cm quadrat. The collected samples were stored in airtight plastic bags in a cool, dark place. On the second day after collection, samples were floated. They were kept individually in buckets containing a volume of water significantly greater than the sample volume for half an hour. Immediately thereafter, the floating material, mainly of vegetable origin, was washed thoroughly in the buckets to remove any remaining snail shells. The washed, snail-free floats were discarded. The material remaining in the dish was washed through a sieve with a mesh size of 0.75 mm. The hole size was chosen so as not to allow the passage of the smallest adult snail specimens. The material remaining on the sieve contained the washed snail shells and other small organic and inorganic debris. This was immediately placed in 70% ethanol and stored.

Later, the snail shells were removed from these samples using a 3.5X magnifying glass. The sorted snail shells were identified to the species level using a stereo microscope (Motic SMZ-168 series, 10X magnification). All determinations were carried out by RF, and they were based on the works of Welter-Schultes (2012), Kerney et al. (1983) and Glöer and Meier-Brook (2003). Only fully developed and intact (not broken or incomplete) shells of live adults were used in the study. Live specimens were considered to be those in which the body of the snail could be observed inside the translucent shell or by looking into the aperture of the specimens. Slugs were not included in the study because this sampling method is not suitable for their quantitative analysis (Cameron and Pokryszko, 2005).

### Statistical analysis

All statistical analyses were performed in the R interactive statistical environment (v4.2.2, R Core Team 2022).

Snail communities were characterised by (i) the total number of specimens, (ii) the number of species and (iii) the normalised Shannon-Weaver diversity: where the diversity measure normalised by the maximum possible diversity, ln R (Shannon and Weaver, 1949). If the different treatments differently affect the changes in snail communities over time one would expect a significant interaction between the variables of treatment (control, mowing and mulching) and sampling time (pre- and post-treatment). To statistically investigate this possibility, we fitted three generalised linear mixed effects models (GLMMs) to our data, one for each community characteristic. We entered treatment, sampling time and their two-way interaction as fixed effects. We also took into account the statistical non-independence caused by spatial proximity by entering random effects into the models. Random effects were entered as sampling point ID nested within parcel ID nested within area ID. Note, however, that we had to remove the random effect terms from the model for Shannon diversity, because this model did not converge as the variance explained by the random effect terms was essentially zero. We used a negative binomial error distribution for the number of specimens, and normal error distributions for the number of species and diversity. Inspecting the model fits with the DHARMa R package (v0.4.6, Hartig 2022) did not indicate any deviations from model assumptions.

We also analysed how treatments affected the number of individuals for the three most common species in the samples (*Carychium minimum*, *Vertigo angustior* and *Vallonia pulchella*). We fitted separate models for each species data with the same fixed and random effects as specified above. We used a negative binomial error distribution for all three species. Model assumptions were also checked by the DHARMa package.

Models were fitted with the ‘glmmTMB’ function of the glmmTMB package (v1.1.7, Brooks et al. 2017). If a significant interaction was detected (by log ratio tests) we tested which treatment groups were significantly different from each other by the emmeans package (v1.8.4-1, Lenth 2023).

## Results

During the study, 5233 living individuals of 20 species were found in the two sample sites combined (**Table 1**). Total number of living individuals in different treatment parcels and periods are given in **Appendix 1**.

**Table 1.** Total number of living individuals in the samples.

| Species | Number of live individuals |
| --- | --- |
| Anisus spirorbis | 2 |
| Carychium minimum | 2031 |
| Carychium tridentatum | 20 |
| Cochlicopa lubrica | 255 |
| Euconulus fulvus | 4 |
| Fruticicola fruticum | 6 |
| Galba truncatula | 2 |
| Laciniaria plicata | 1 |
| Nesovitrea hammonis | 20 |
| Oxyloma elegans | 5 |
| Pseudotrichia rubiginosa | 60 |
| Punctum pygmaeum | 8 |
| Succinella oblonga | 39 |
| Trochulus hispidus | 12 |
| Vallonia pulchella | 551 |
| Vertigo angustior | 2021 |
| Vertigo antivertigo | 152 |
| Vertigo pygmaea | 4 |
| Vitrina pellucida | 1 |
| Zonitoides nitidus | 39 |

Before treatment, we found no significant differences between the mowed, mulched and control parcels in either the number of live individuals, number of species or Shannon diversity values (**Appendix 2**).

Looking next at the effects of treatment, we found a significant interaction between treatment and sampling time for all community characteristics: number of living individuals (χ _2_^2^ = 6.99, p = 0.03), species number (χ _2_^2^ = 11.32, p = 0.01) and Shannon diversity (χ _2_^2^ = 6.26, p = 0.04). This indicates that different treatments affected the snail communities differently.

Next, we examined how snail populations changed due to each treatment (**Appendix 3**, **Fig 3**).

**Figure 3.**
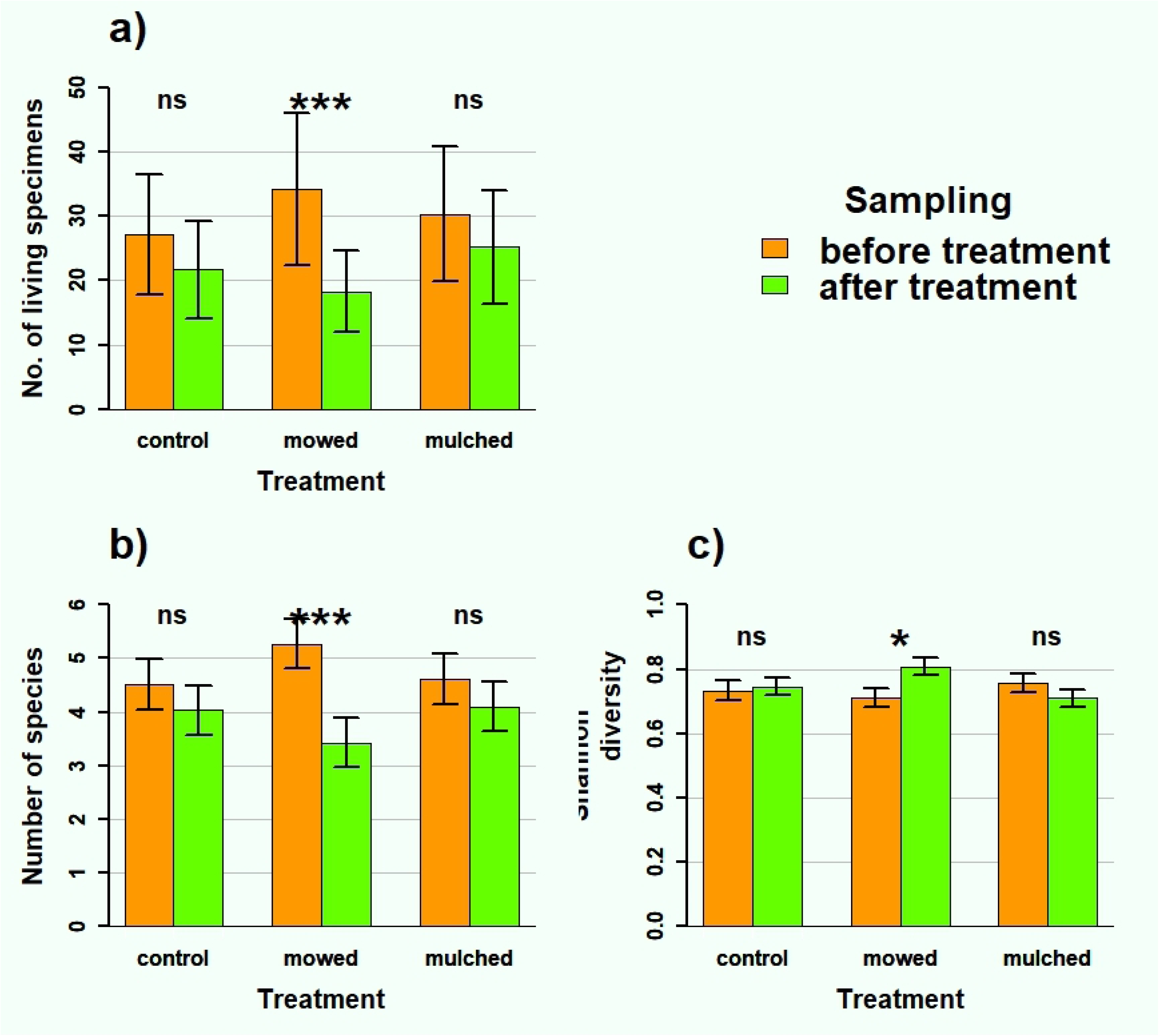
The impact of treatment The impact of treatment a) on the number of living individuals, b) on the number of species and c) on diversity. Vertical error bars indicate standard errors of the estimates derived from the GLMM models (***: p<0.001, *: p<0.05, ns: p>0.05).

The estimated number of living individuals decreased in all three management types, but this decrease was only significant in the mowed areas, indicating that the mowing itself likely caused a strong decrease. In terms of species numbers, the treatment also caused a significant decrease in the mowed areas. For diversity, we also found a difference only in the mowed areas, but in this case diversity increased as a result of mowing.

In addition to community-level effects, we investigated whether treatments had species-level effects on the three most common species (*Carychium minimum*, *Vallonia pulchella*, *Vertigo angustior*). In *Carychium minimum* the abundance decreased significantly with sampling time (χ21 = 97.16, p < 0.001) independently of treatment (effect of treatment: χ22 = 0.72, p = 0.696; interaction: χ22 = 2.85, p = 0.241). In the case of *Vallonia pulchella* none of the effect of treatments (χ^2^_2_ = 3.55, p = 0.169) and sampling times (χ^2^_1_ = 2.62, p = 0.105) neither of their interaction (χ^2^_2_ = 3.98, p = 0.137) was significant. Only in the case of *Vertigo angustior*, we found that treatments (interaction: χ^2^_2_ = 7.90, p = 0.019) differently affect the change in number of individuals; mulching (posthoc test: z = −3.25, p = 0.001) significantly increased the number of individuals while the other two treatments did not have such an effect (control, posthoc test: z = −1.53, p = 0.127; mowing, posthoc test: z = −0.81, p = 0.417; **Fig 4**).

**Figure 4.**
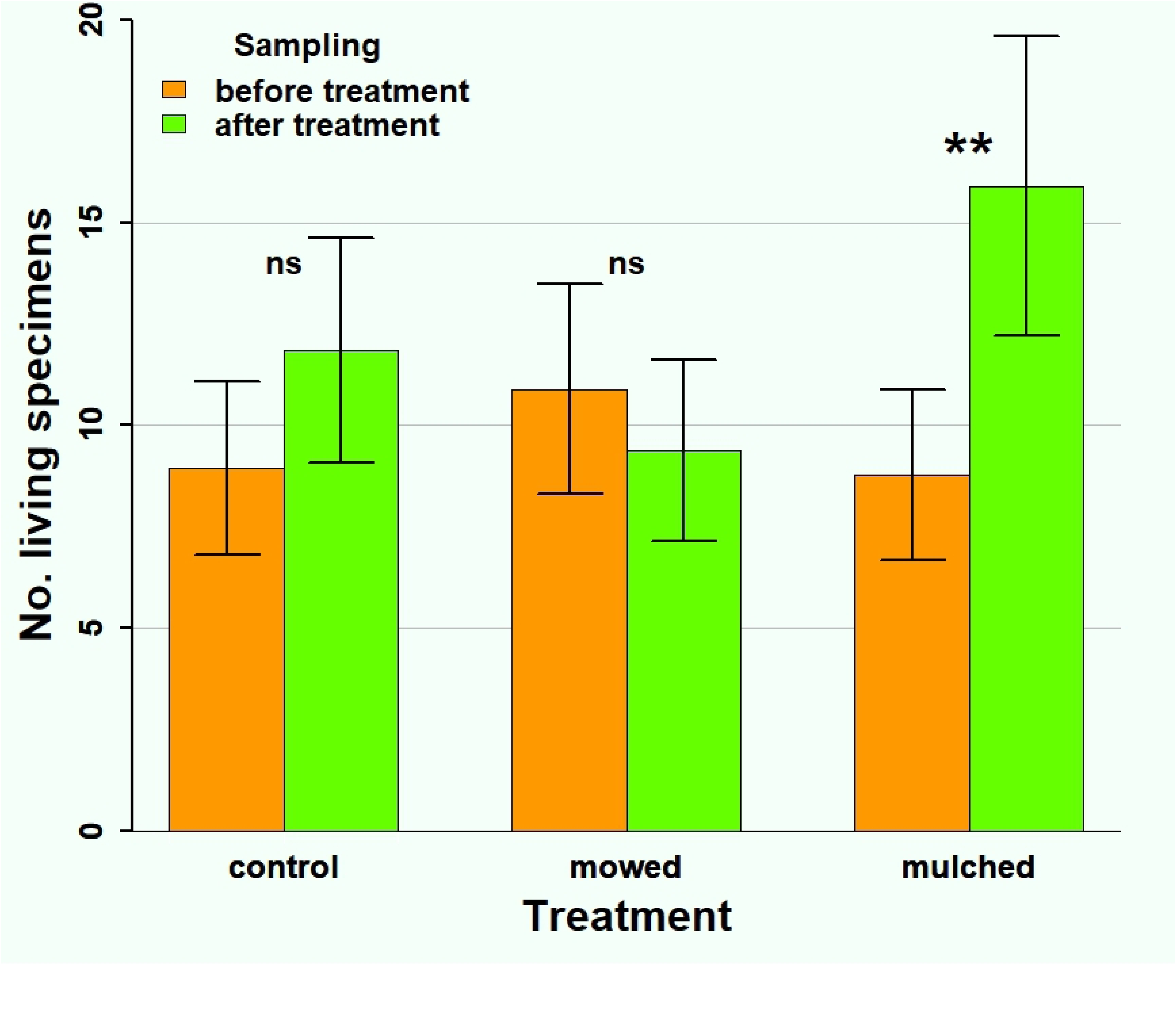
The impact of treatment on the number of living *Vertigo angustior* individuals. Vertical error bars indicate standard errors of the estimates (**: p<0.01, ns: p>0.05).

## Discussion

We found that different treatments had different impacts on the snail communities. In the mowed areas, negative changes in the snail community were observed one year after the treatments. Significant declines in both live individuals and species richness indicate that mowing had a detectable negative impact on snail communities even after one year. In comparison, mulched and control areas experienced no significant changes.

Our finding that diversity increased on mowed plots might seem to contradict our claim that mowing can have a negative effect on snail communities. Under certain conditions, however, diversity index can increase despite a decrease in number of individuals and species. For instance, if more the decrease is larger in more abundant species then distribution of individuals among species becomes more even and hence the diversity index increses.

There has been relatively little research on the effects of mowing on snails, and, as far as we know, there is no previous work on the effects of mulching on them. Previous malacological research on mowing has mainly focused on several years long effects. For instance, Pech and colleagues (2015; working in oligotrophic grasslands), and Farkas and colleagues (2024; on valley-bottom wet meadows) found long-term reductions in abundance and species numbers due to mowing, but Wehner and colleagues (2019; over a wide range of meadows) found an increase in snail community diversity in mowed areas, while no difference in abundance was found. All three studies are similar in that they were conducted on areas that were previously mowed (albeit for varying durations), while our current study was conducted in wet meadows that were previously untreated for decades. In addition, the above studies investigated the effects of mowing not by comparing samples taken before and after treatment on the same site, but by analyzing data from mowed and adjacent unmowed areas. Despite these methodological differences, our result on the detrimental effects of mowing is similar to two of these previous studies.

Our findings pertain to snail communities, yet wet meadows harbor numerous other abundant invertebrate taxa, which may respond differently to different management practices. Mowing had negative effects on spiders (Cattin et al. 2003), orthopterans (Cizek et al. 2012, Chisté et al. 2016) and dung beetles (Frank et al. 2017). Other studies have found negative effects of mulching on moth (Georgi et al. 2023) and wasp (Georgi et al. 2022) communities. There are studies that conclude that mulching is less beneficial than mowing for earthworms, oribatids (Pizl and Stary 2001), and butterflies (Schmitt 2003). However, the number of studied invertebrate groups is limited. This makes it difficult to generalize about the effects of treatments on entire invertebrate communities of a habitat. Moreover, the range of habitats studied is broad, with varying initial conditions: some studies monitor the effects of different ongoing treatments (Chisté et al. 2016, Frank et al. 2017, Schmitt 2003), while others established new experimental setups on habitats that have earlier undergone some form of regular treatment (Georgi et al. 2022, 2023). However, there are no previous studies where the initial state of the habitat, like ours, was untreated for decades. Going forward, significantly more research is needed to investigate the effects of grassland management on invertebrates in order to ensure that different habitats receive the most suitable management practices for the widest possible range of taxa.

Our results indicate that members of snail communities did not respond uniformly to the management. Of the three most common species we examined, only *Vertigo angustior* showed a treatment specific response. Both *Carychium minimum* and *Vertigo angustior* are typical of wetland habitats, but only the latter was affected by the management. *Vallonia pulchella* has a wider range of habitat preferences, and was unaffected by the treatment.

Similar results were observed in management studies with other taxa. Cizek and colleagues (2012) found that the abundance of different ground beetle species changed differently with the same treatment. Pižl and Starý (2001) showed differential changes in abundance for different species of oribatids living above the soil surface in response to management. Both sources agree that generalist species tolerate changes better than specialists. *Vertigo angustior* is still considered a common species in Hungary, but its distribution in Western Europe has drastically declined, which can be attributed to changing and increasingly intensive agriculture (European Environment Agency 2013). Therefore, the appropriate choice of management practices is of paramount importance in its case.

Our study was designed to examine the one year effects of a single intervention. Negative effects of mowing were already detectable one year after treatment. We assumed that no significant changes in the chemical and soil properties of the site could have occurred in such a short time. The one year negative effects on snail communities at both sites for all treatments might have been mainly due to effects such as sudden increase in irradiance, increases in temperature, decreases in humidity and organic matter.

Management practices usually have an effect due to changes in habitat characteristics (Wehner et al. 2019). Over a longer period of time, habitat characteristics to which snail or other invertebrate communities were originally adapted can change significantly. Sensitive species may recede, while others, adapting to new opportunities, may advance. Careful selection of management conditions can significantly reduce their negative impact on invertebrates. These include staggering managements throughout the year or leaving parts of the habitat unmanaged (Pech et al. 2005, Cizek et al. 2012,). The negative effects of treatments could also be prevented or reduced by tailoring them to the life cycle of invertebrates (Georgi et al. 2022, 2023). It may also be beneficial not to manage an area every year, but this can threaten the survival of the whole habitat after a certain reduction in intensity (Ryser et al. 1995, Moog et al. 2002).

## Conclusions

Wet meadows have been sites of grassland management since ancient times. The evolution of methods and increasingly intensive agricultural practices are significantly transforming these valuable habitats. While the impact of treatments on vegetation is somewhat understood, little is known about the fate of animal communities, especially invertebrates. Through a controlled grassland management experiment, we examined the effects of mowing and mulching on the snail communities of wet meadows. Our results indicate that significant changes are observed just one year after initiating treatment. The negative impact of mowing was clearly evident within this time frame, while mulching did not lead to significant changes. Mulching thus seems to be a practical way for short-term interventions of grassland areas without the negative effects of mowing. Sustaining grasslands in the long term requires attention not only to habitat maintenance but also to preserving their quality and biodiversity. Therefore, selecting appropriate methods is crucial. Within a given area, numerous taxa are present, each responding differently to interventions. Hence, conducting similar studies across many and diverse invertebrate groups is necessary. In addition, whilst one year conclusions regarding the effects of different management methods are useful, there is still a lack of long-term monitoring of grassland management, particularly regarding mulching. Initiating, supporting, and maintaining more such long-term research projects is crucial. In addition to economic considerations, ecological aspects must also be taken into account to ensure sustainability for the future.

## Acknowledgement

We are grateful to Tamás Székely, Jr. for his help to correct our English.

## Appendix

**Appendix 1.**
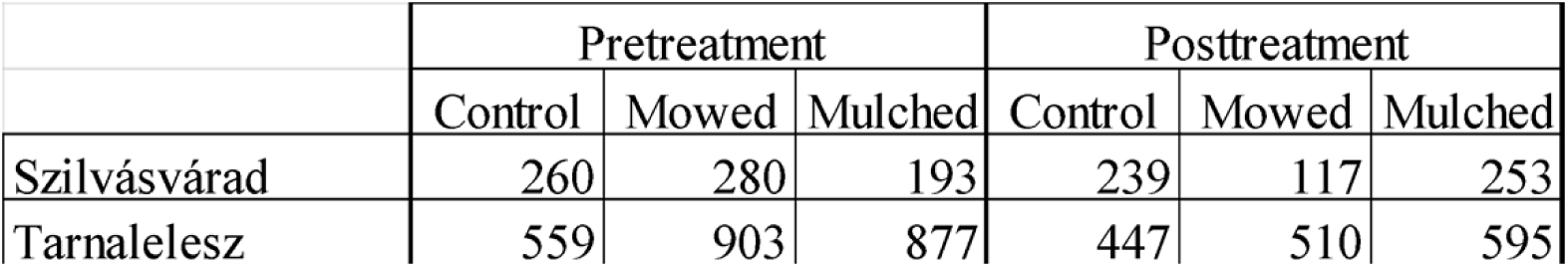
Total number of living individuals in different treatment parcels and periods in the two study area.

**Appendix 2.**
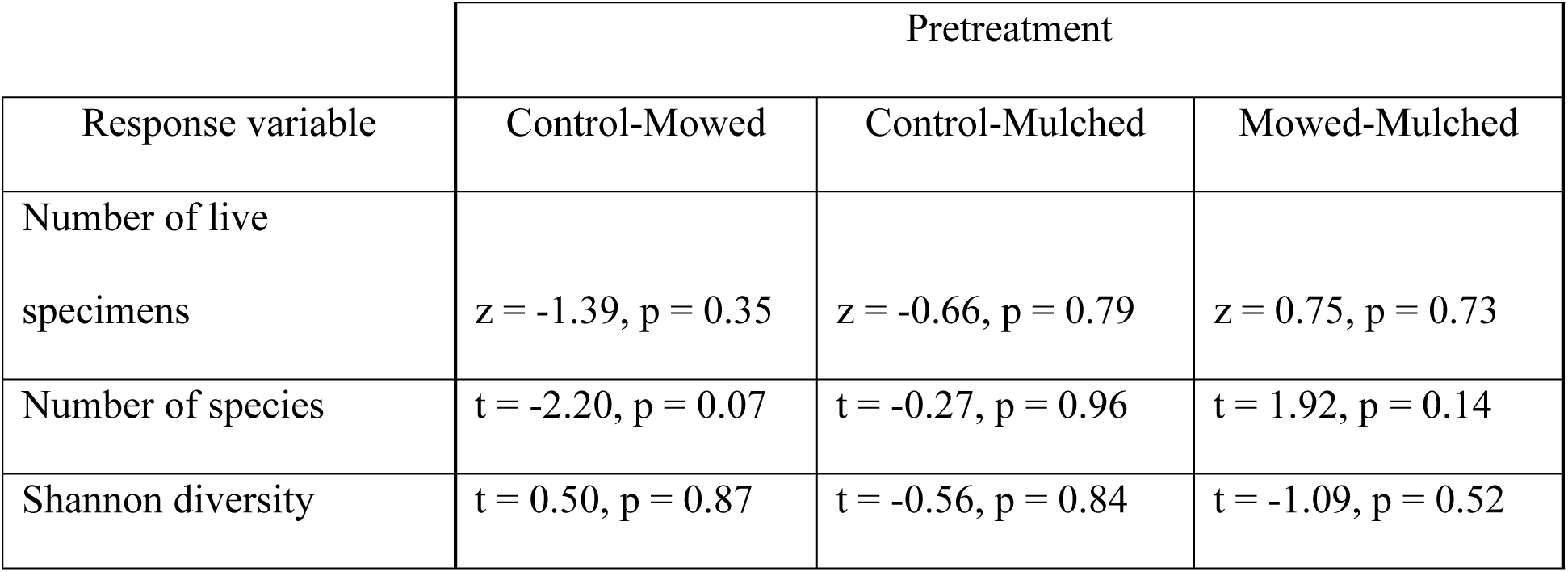
Pairwise comparison of treatments before the treatments. Posthoc tests from GLMMs separately fitted to each response variable (see the main text for details).

**Appendix 3.** Changes occurring in different treatment parcels due to treatment. Posthoc tests from GLMMs separately fitted to each response variable (see the main text for details). Bold font indicates significant differences.

| Response variable | Control | Mowed | Mulched |
| --- | --- | --- | --- |
| Number of live specimens (pre-/post-treatment) | $z = 1.62, p = 0.10$ | **$z = 4.68, p < 0.0001$** | $z = 1.43, p = 0.15$ |
| Number of species (pre-/post-treatment) | $t = 1.53, p = 0.13$ | **$t = 5.86, p < 0.0001$** | $t = 1.62, p = 0.11$ |
| Shannon diversity (pre-/post-treatment) | $t = -0.31, p = 0.76$ | **$t = -2.39, p = 0.02$** | $t = 1.17, p = 0.24$ |

